# Genome-wide association study identifies 44 independent genomic loci for self-reported adult hearing difficulty in the UK Biobank cohort

**DOI:** 10.1101/549071

**Authors:** Helena RR. Wells, Maxim B. Freidin, Fatin N. Zainul Abidin, Antony Payton, Piers Dawes, Kevin J. Munro, Cynthia C. Morton, David R. Moore, Sally J Dawson, Frances MK. Williams

## Abstract

Age-related hearing impairment (ARHI) is the most common sensory impairment in the aging population; a third of individuals are affected by disabling hearing loss by the age of 65^1^. ARHI is a multifactorial condition caused by both genetic and environmental factors, with estimates of heritability between 35% and 55%^2–4^. The genetic risk factors and underlying biological pathology of ARHI are largely unknown, meaning that targets for new therapies remain elusive. We performed genome-wide association studies (GWAS) for two self-reported hearing phenotypes, hearing difficulty (*HDiff*) and hearing aid use (*HAid*), using over 250,000 UK Biobank^5^ volunteers aged between 40-69 years. We identified 44 independent genome-wide significant loci (P<5E-08), 33 of which have not previously been associated with any form of hearing loss. Gene sets from these loci are enriched in auditory processes such as synaptic activities, nervous system processes, inner ear morphology and cognition. Immunohistochemistry for protein localisation in adult mouse cochlea indicate metabolic, sensory and neuronal functions for *NID2, CLRN2* and *ARHGEF28* identified in the GWAS. These results provide new insight into the genetic landscape underlying susceptibility to ARHI.

---

ARHI is characterised by a non-syndromic bilateral, sensorineural hearing loss that progresses with increasing age and is an established risk factor for depression^6–8^ and dementia^9–12^. Hearing loss was ranked fourth in the latest study into the Global Burden of Diseases^13^, yet hearing amplification devices are the only treatment option currently available for ARHI. ARHI is expected to be a highly genetically heterogeneous trait given that over 150 genetic loci have been identified in non-syndromic hereditary hearing loss alone (https://hereditaryhearingloss.org/). Previous GWAS of ARHI have identified a small number of promising candidate genes, though there has been poor replication of findings to date, possibly reflecting varied phenotyping approaches and limited sample sizes^14–24^.

We conducted two GWAS using the self-reported hearing difficulty and hearing aid use of UK Biobank (UKBB) participants and refined our results using a combination of conditional analysis, replication analysis, *in silico* annotation and *in vivo* expression analysis (see Figure 1 for study design). Our aim was to identify the genetic components of adult hearing impairment in the UK population and provide insight into the pathology of ARHI.

**Figure 1.**
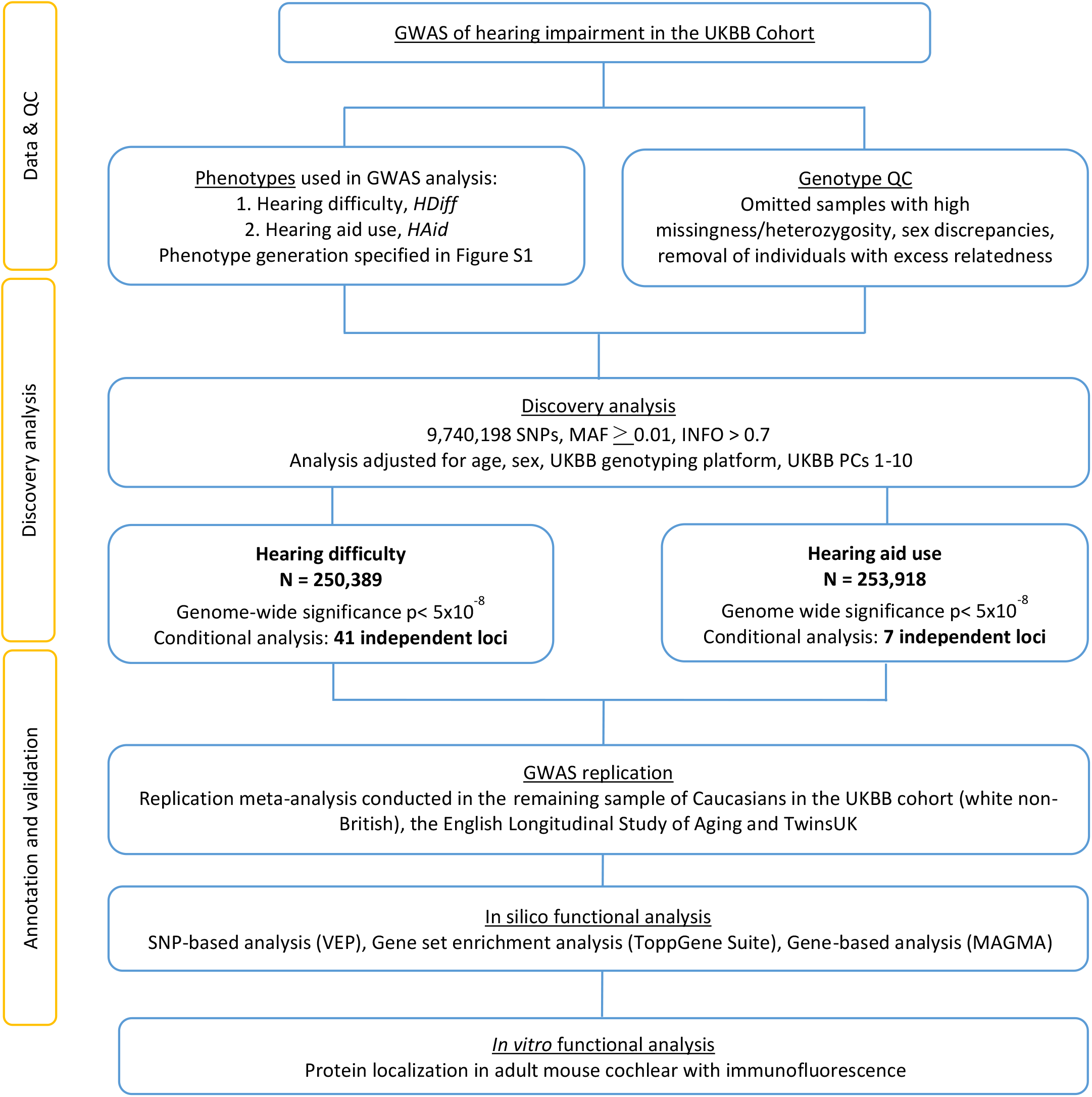
Workflow schematic for discovery and validation of associated loci. N, sample size; QC, quality control; PC, principal components; MAF, minor allele frequency; INFO, quality metric, combination of imputation score and dosage confidence

UKBB participants were categorised using a case-control design based on responses to questions regarding hearing difficulty (*HDiff*, n=498,281) and hearing aid use (*HAid*, n=316,629) (Supplementary Figure 1). A linear mixed-effects model was used to test for association between 9,740,198 SNPs and the two traits, using BOLT-LMM v.2^25^, which corrects for population stratification and within sample relatedness. Following additional quality control filters and selection of white British participants (described in online methods), the final samples for association analyses were n=250,389 for *HDiff* and n=253,918 for *HAid* (Supplementary Figure 1).

The studies identified 2,080 and 240 SNPs at genome-wide significance (P<5E-08) for *HDiff* and *HAid* analysis, respectively (Figure 2 and Supplementary Figure 2). Conditional and joint analysis using GCTA-COJO^26^ identified 41 and seven independent loci associated with *HDiff* and *HAid*, respectively, resulting in 44 independent loci when accounting for common overlap between the two phenotypes. SNP heritability estimates for the two traits calculated with BOLT-LMM (h2g) were 0.117 +/-0.001 for *HDiff* and 0.029, +/-.001 for *HAid*. Estimates recalculated to the liability scale are 0.19 and 0.13 for *HDiff* and *HAid* respectively.

**Figure 2.**
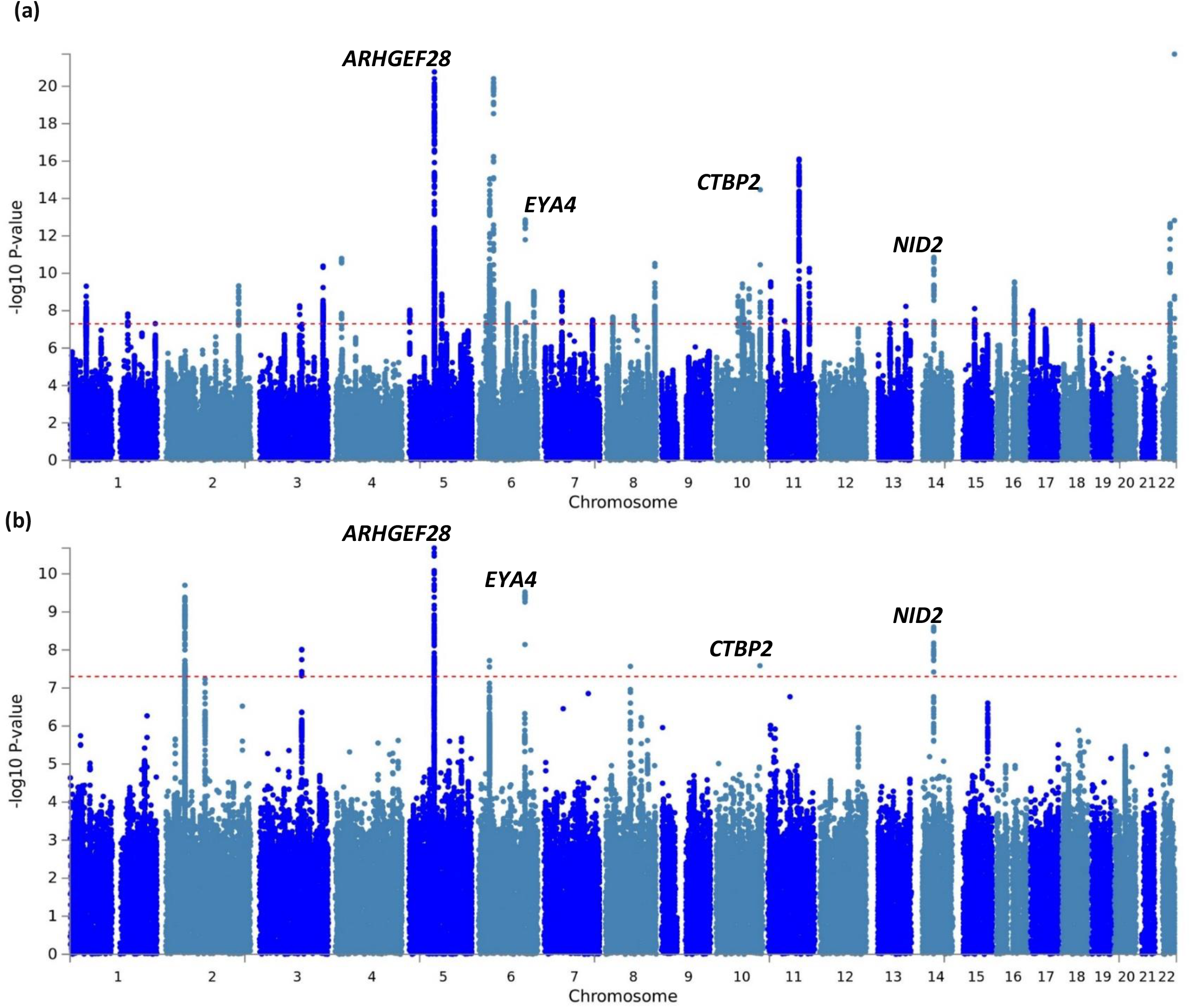
Manhattan plots displaying GWAS results for (*a*) Hearing difficulty, and (*b*) Hearing aid use phenotypes. The Manhattan plots display the P values of all SNPs tested in discovery analysis. The threshold for genome wide significance (p<5×10^−8^) is indicated by a red dotted line. Loci that reached genome-wide significance in both phenotypes are annotated with gene symbol.

The Variant Effect Predictor (VEP)^27^ was used to map independent lead SNPs to the nearest protein coding genes, using the GRCh37 genomic reference. Of 41 independent SNPs associated with *HDiff*, six variants lie in exons, four of which result in missense mutations in *EYA4, CDH23, KLHDC7B* and *TRIOBP,* 21 SNPs lie within introns and 14 are intergenic (Table 1). Six of the independent SNPs associated with *HAid* reside in intronic regions and 1 is intergenic. Significant gene loci common to both traits were *NID2, ARHGEF28, CTBP2* and *EYA4* (Supplementary Figure 3). Variants within *EYA4* have been reported in autosomal dominant non-syndromic hearing loss^28–30^, while *NID2* and *ARHGEF28* are new associations with hearing impairment. *CTBP2*, though not previously linked to genetic risk of ARHI, encodes a protein component of the inner ear hair cell pre-synaptic ribbon^31^.

**Table 1.**
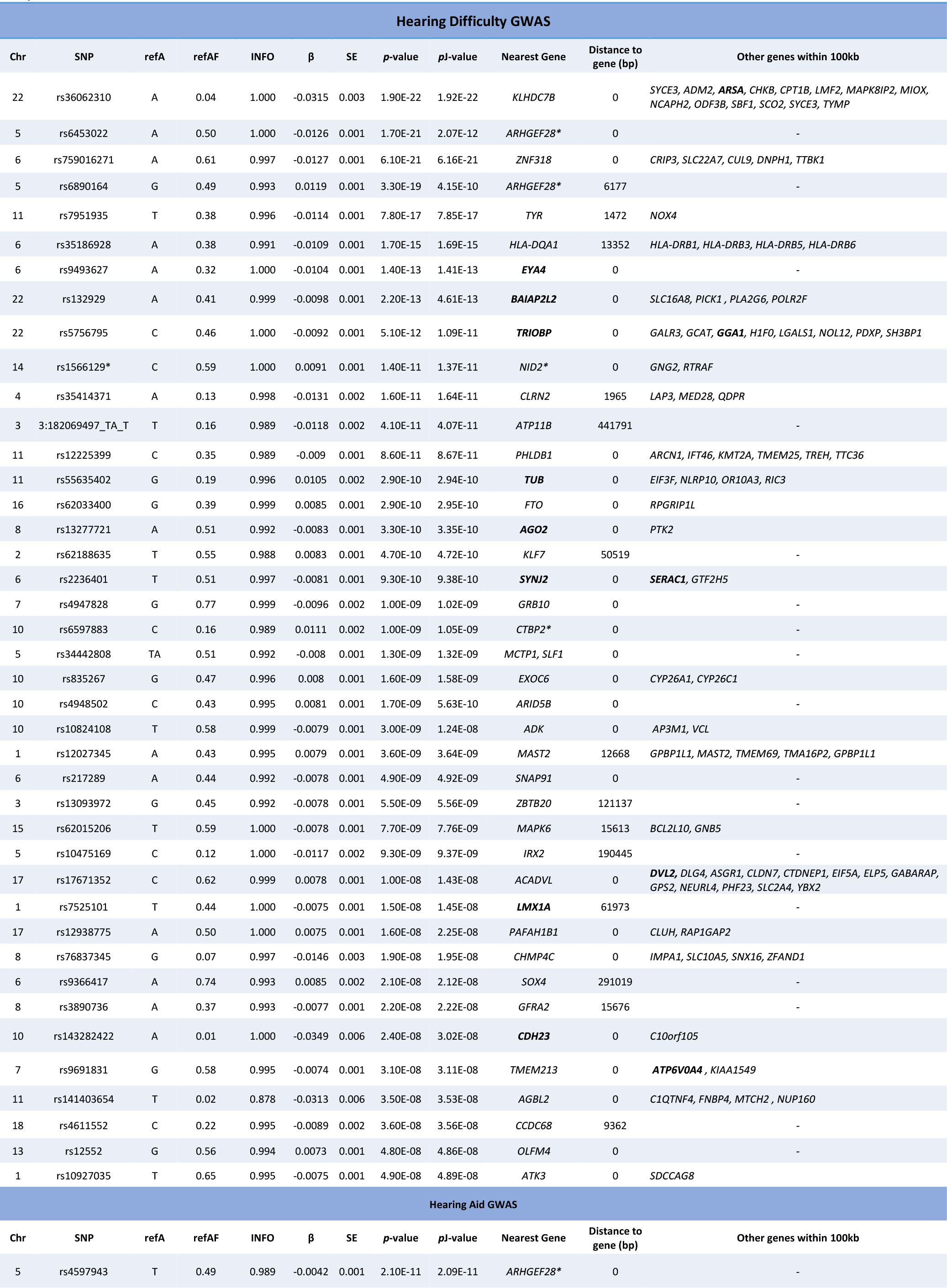
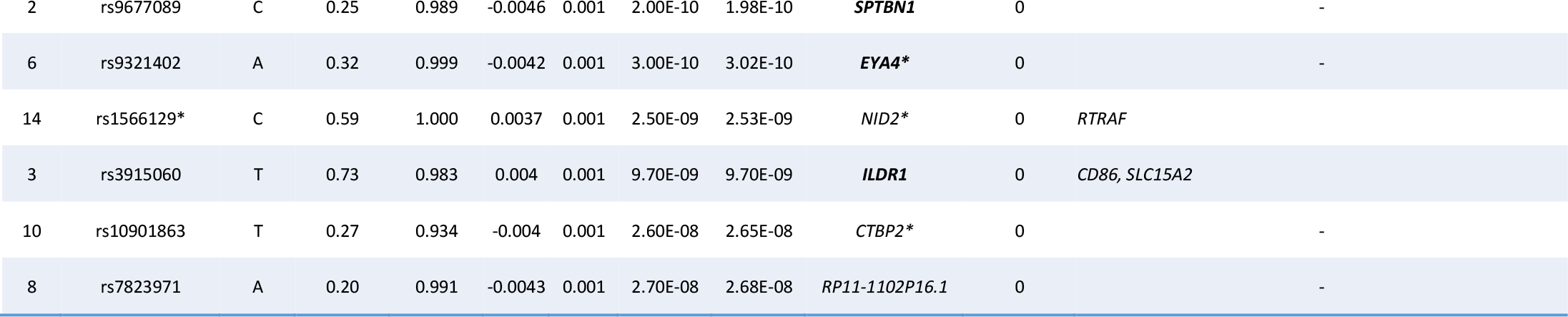
Independent SNPs significantly associated (P<5 × 10–8) with the two phenotypes regarding hearing ability in the UK Biobank discovery sample. Results output from BOLT-LMM and GCTA-COJO. Chr., chromosome; SNP, single nucleotide polymorphism; refA, reference allele in COJO-GCTA analysis; refAF, frequency of effect allele in COJO-GCTA analysis sample; INFO, quality metric, combination of imputation score and dosage confidence; β, effect size from BOLT-LMM approximation to infinitesimal mixed model; SE, standard error of the effect size; *p*-value, infinitesimal mixed-effects model association test p-value; *p*J-value, p-value from a joint analysis of all the selected SNPs; Nearest Gene, protein-coding gene in closest proximity to SNP; Distance to gene (bp), distance in base pairs between SNP and nearest gene, a value of 0 indicates the SNP lies within the gene; Other genes within 100kb, list of genes within 100kb of the SNP. Bold font denotes genes previously linked to hearing phenotypes in mice or humans, * denotes SNP or gene common to both *HAid* and *HDiff* studies.

**Figure 3.**
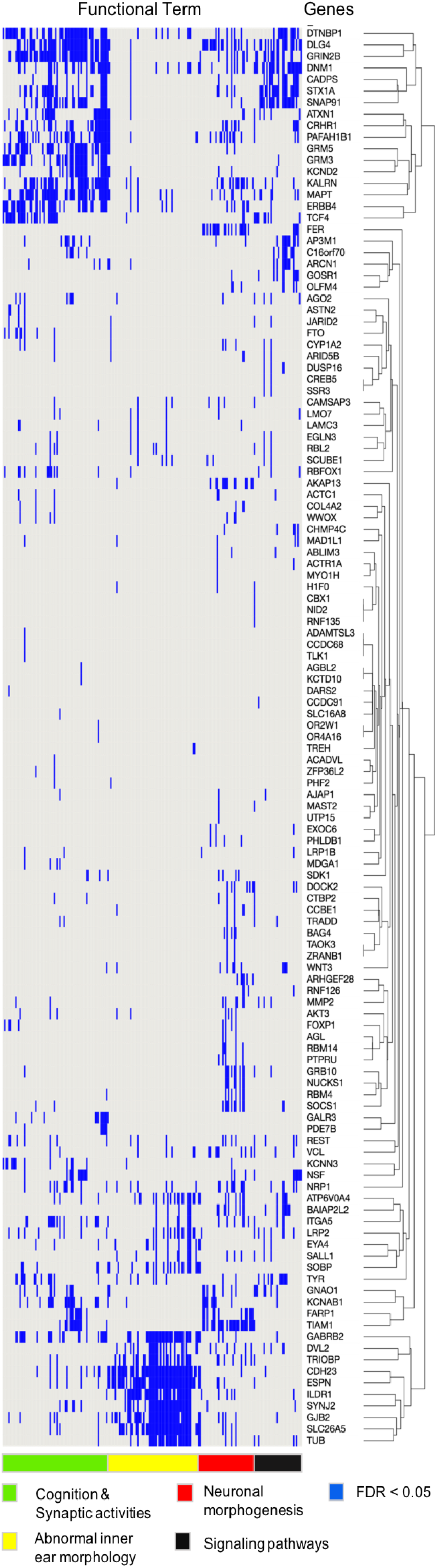
Heatmap of the enriched functional terms for genes mapped to lead SNP at suggestive level (*HDiff* analysis), using ToppGene Suite. Genes for enriched functional terms at FDR 0.05 are in blue. Genes and terms were grouped using clustering of presence and absence status of genes in respective functional terms. Functional terms include GO Biological Process, GO Molecular Function, GO Cellular Component, Mouse Phenotype, Pathway, and Disease.

Replication was attempted for the lead SNPs (41 *HDiff* and 7 *HAid*) by meta-analysing three independent samples; the remaining Caucasians in the UKBB cohort (white, non-British Europeans), TwinsUK, and the English Longitudinal Study of Aging (ELSA), totalling *HDiff* N = 30,765 and *HAid* N = 35,004 (see online methods). Two SNPs in *ZNF318* and *NID2* reached significance in the *HDiff* replication analysis (Bonferroni correction 0.05/41=0.0012, P<0.0012), and one SNP in *ARHGEF28* replicated in *HAid* analysis at the significance threshold (0.05/7=0.00714, P<0.00714). An additional 14 SNPs reached nominal significance (Supplementary Tables 1 and 2).

We investigated whether any of the candidate genes identified in adult hearing in previously published genetic association studies were replicated within the discovery White British sample (Table 2) and found two previous variant associations located in close proximity to *ISG20* and within *TRIOBP,* which were identified in a GWAS performed with data from electronic health records^24^.

**Table 2.**
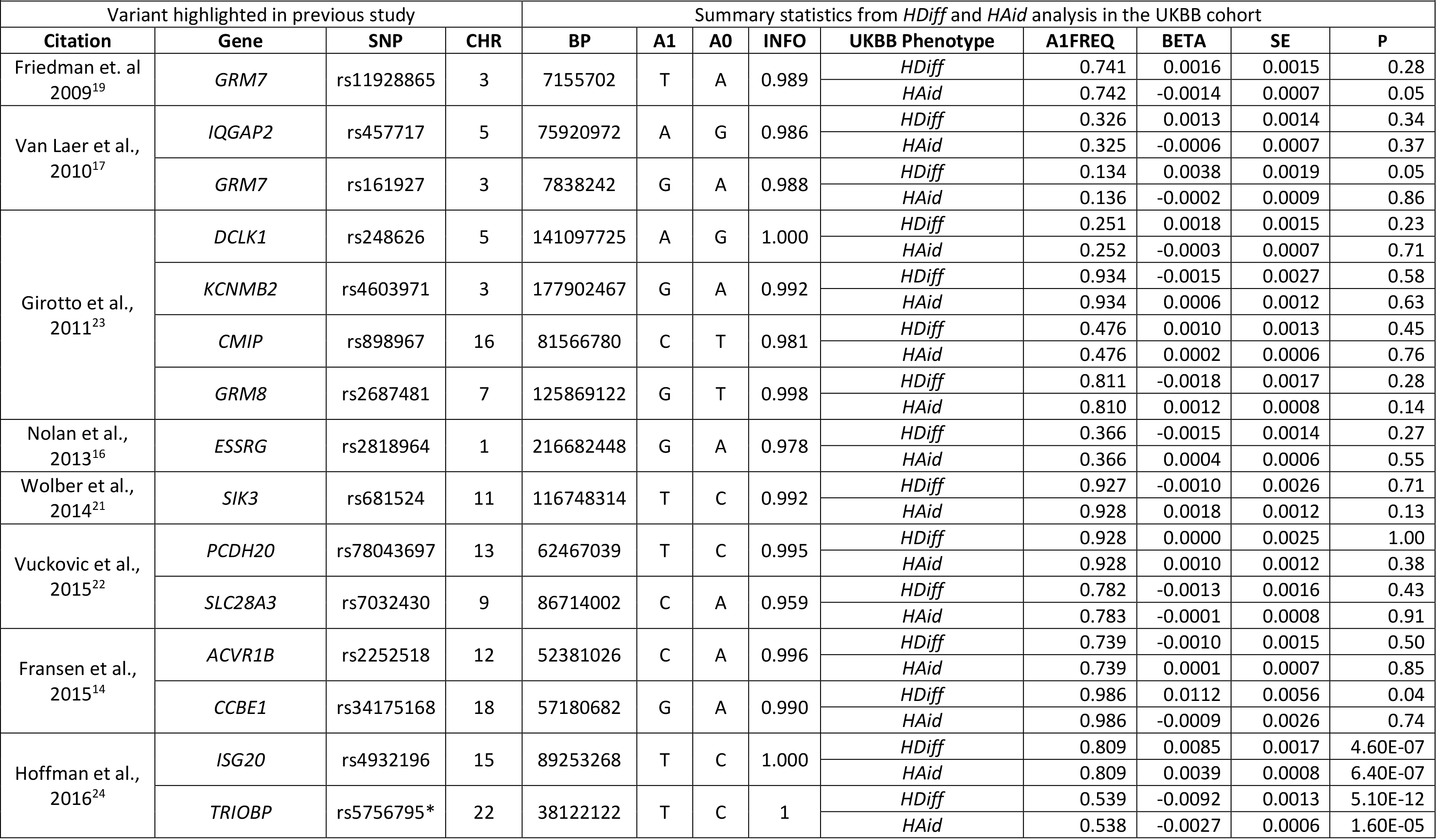
Summary statistics from *HDiff* and *HAid* GWAS analysis, at SNPs highlighted in previous adult hearing loss GWAS. Study, publication of previous finding; Gene, gene highlighted in the referenced publication as the lead SNP is either located in the gene region or in close proximity; SNP, single nucleotide polymorphism; CHR, Chromosome; BP, base position; A1, effect allele in analysis; A0, reference allele; INFO, quality metric, combination of imputation score and dosage confidence; UKBB phenotype, phenotype used in this study; A1FREQ, frequency of effect allele in analysis sample; BETA, effect size from BOLT-LMM approximation to infinitesimal mixed model; SE, standard error of the effect size; p-value, infinitesimal mixed model association test p-value. This study did not analyse SNP rs58389158, but analysed rs5756795 which is in complete LD with this SNP in the British population, and referenced in the previous study. This is denoted by * in the table.

While *ISG20* is a novel association, mutations in *TRIOBP* cause one form of autosomal recessive non-syndromic deafness, DFNB28^32,33^. No other lead variants from previous ARHI genetic studies were replicated at nominal level in our analysis, including the first reported ARHI associated gene variant in *GRM7*^15^.

Functional gene annotation was undertaken with genes mapped from SNPs associated at a suggestive level in the *HDiff* association analysis. Genes were significantly enriched in a number of processes required for auditory function: synaptic activities, trans-synaptic signalling, nervous system processes, modulation of chemical synaptic transmission, positive dendritic spine morphogenesis, and inner ear morphology as well as cognition, learning or memory. These genes were also significantly enriched with mouse phenotype ontologies, mostly relating to inner ear abnormalities and abnormal auditory brainstem response, and were significant at FDR 0.05 (Figure 3). As well as suggesting pathogenic pathways, this finding demonstrates the shared genetic pathology in mouse and human auditory systems, supporting the use of mouse models to study human auditory function.

*In silico* tissue-specific gene expression analysis undertaken with MAGMA^34^ indicates a significant association between *HDiff* suggestive genes and transcription levels of genes in brain (P = 5.4E-04; Supplementary Figure 4). This finding could be due to the fact that sensory cells of the inner ear are of neural origin and a substantial amount of neuronal tissue expression data is available in comparison to the limited datasets derived from cochlear tissue.

We investigated expression of putative novel hearing genes *NID2, ARHGEF28* and *CLRN2* in adult mouse cochlea using immunohistochemistry. The lead SNP in *NID2* in both *HDiff* and *HAid* is located in intron 5 and replicated in the *HDiff* meta-analysis. Two independent lead SNPs were identified at the *ARHGEF28* locus in the *HDiff* analysis, along with a third SNP in the *HAid* analysis which replicated in the meta-analysis. The lead independent SNP at the *CLRN2* locus in the *HDiff* analysis is within 2kb of *CLRN2*, although several other genes are within 100kb. Because *CLRN1,* a paralog of *CLRN2,* is expressed in hair cells and mutations in *CLRN1* cause autosomal recessive Usher syndrome Type-3 with progressive sensorineural hearing loss,^35,36^ we investigated whether clarin-2 is also expressed in the inner ear.

Immunostaining for nidogen-2, a basement membrane component encoded by *NID2*, was most prominent in the epithelial lining of the inner spiral sulcus between the tectorial membrane and the inner hair cell (Figure 4, Supplementary Figure 5), as well as localizing to nerve fibres and blood vessel basement membranes, as has been noted in other tissues previously ^37^.

**Figure 4.**
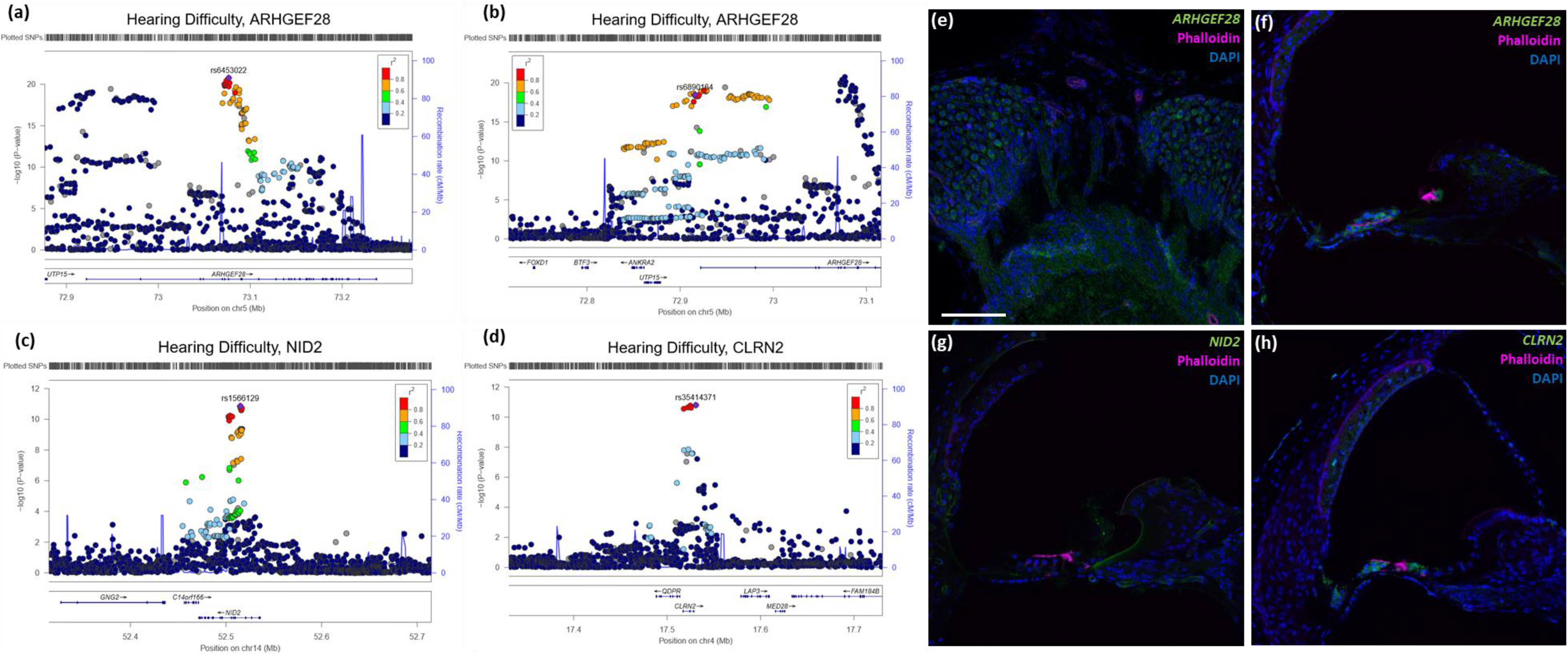
Cochlear expression of three putative hearing genes identified in *HDiff* and *HAid* GWAS. **(a,b,c,d)** Locus zoom plots of associated loci, generated with *HDiff* summary statistics. Four associated loci are plotted which have lead SNPs in or in proximity to *ARHGEF28* **(a,b)**, *NID2* **(c)**, and *CLRN2* **(d)**. Purple indicates lead independent SNP generated from GCTA-COJO conditional analysis. Colouring of remaining SNPs is based on linkage disequilibrium (LD) with the lead SNP. The genes within the region are annotated, and the direction of the transcripts is shown by arrows. Two independent regions were identified within the *ARHGEF28* locus; both are shown. **(e,f,g,h)** Immunofluorescence images of adult mouse cochlea, spiral ganglion neurons **(e)** and organ of Corti **(f-h)**. Vibratome sections stained with the three proteins of interest in mouse inner ear; DAPI (blue) and Phalloidin (magenta) were also used for staining of actin and nuclei respectively. **(e)** Anti-ARHGEF28 staining is observed in the neuronal cell bodies and axons. **(f)** Anti-ARHGEF28 (green) is mainly observed in outer and inner hair cells. **(g)** Anti-NID2 (green) staining is observed lining blood vessels and the epithelial lining of the inner spiral sulcus. **(h)** Anti-CLRN2 (green) staining is observed in outer and inner hair cells, in addition to the stria vascularis. The scale bar in image **(e)** represents 100µm. The scale is consistent for all images in this figure.

Similar to clarin-1, clarin-2 immunostaining localised to the inner and outer hair cells, the primary sensory cells of sound detection, suggesting it may also be necessary for hearing (Figure 4, Supplementary Figure 5).

*ARHGEF28* encodes Rho Guanine Nucleotide Exchange Factor 28, for which immunostaining was observed in both hair cells and the spiral ganglion neuron cell bodies and axons (Figure 4, Supplementary Figure 5). Previous reports demonstrate a role for *ARHGEF28* in regulation of neurofilaments^38,39^ and axon growth and branching^40^. It has also been implicated in the pathogenesis of motor neuron disease through formation of neurofilament and *ARHGEF28* aggregates^41^.

Our study should be received in the context of its limitations; first, there is currently a lack of adequately powered studies with which to replicate our results. Despite meta-analysing three cohorts, the replication sample remains an order of magnitude smaller than the discovery set. However, the identification of known hearing genes, gene annotation analysis and the results of *in vivo* expression provide support and putative mechanisms for involvement of these genes in hearing loss.

Second, we cannot confirm age at hearing difficulty onset or hearing aid prescription, making an accurate diagnosis of ARHI a challenge. Some of the associations, for example, may be driven by the presence of individuals with congenital hearing impairment due to highly penetrant variants. We reduced the likelihood of this by implementing a minor allele frequency (MAF) cut-off of 0.01 (i.e., higher than the rate of variants in congenital deafness) and by excluding participants who selected ‘I am completely deaf’ in the UKBB questionnaire.

In summary, we have conducted the largest GWAS to date on adult hearing and have identified 44 associated independent loci. Although several genes identified are known to have a role in congenital deafness or have been identified in mouse models, 33 of the 44 loci identified have not previously been associated with hearing loss phenotypes in humans or mice. For three such genes we demonstrated localised cell specific expression within the mouse adult cochlea. This study demonstrates that self-reported hearing loss in adults is suitable for use in association studies using large cohorts such as the UKBB. Our results present a framework for further study into the auditory pathways influenced by the genomic loci identified.

## Online methods

### Participants

The cohort used for discovery association analysis consisted of UK Biobank (UKBB) participants with ‘White British’ ancestry. The UKBB sample classification ‘White British’ is derived from both Principal Component (PC) analysis and self-declared ethnicity^42^. Samples with excess heterozygosity, excess relatedness and sex discrepancies were identified and removed prior to analysis, resulting in samples sizes of n =250,389 and n = 253,918 for hearing difficulty (*HDiff*) and hearing aid (*HAid*) use respectively.

For replication analysis, we used the UKBB ethnic group ‘Caucasians’ (white non-British Europeans). To assign participants into discrete ancestry clusters, we used the 1st and 2nd PC vectors provided by UKBB. A k-means clustering algorithm was applied to generate clusters for each PC. We then combined cluster indices for the PCs (1.1, 1.2, …, 5.5), compared them against self-reported ancestry and assigned the ancestry group accordingly. If contradictory, the pairwise clusters took precedence over the self-report grouping.

The two other samples used for replication analysis were the English Longitudinal Study of Aging (ELSA) and TwinsUK. These datasets were selected as they consist of predominantly Caucasian samples and include relevant questionnaire data. ELSA is a longitudinal study, consisting of around 12,000 respondents from the Health Survey for England. Eight waves of data collection have been completed since 2002^43^. TwinsUK is the largest adult twin registry in the UK and comprises over 13,000 healthy twin volunteers aged 16-98. Collection of data and biologic materials commenced in 1992 and is ongoing. During study participation, twins complete health and lifestyle questionnaires and attend clinical evaluations^44^.

### Phenotype definitions

Two phenotypes were derived for this study; a phenotype representing self-reported hearing difficulty (*HDiff*) and a phenotype representing self-reported hearing aid use (*HAid*). Participants in the UKBB study completed a touchscreen questionnaire during their visit to the assessment centre, which included questions regarding hearing status. Participants were assigned case/control status based on their responses to questionnaire measures regarding hearing difficulty and hearing aid use. Details of how the UKBB phenotype was derived are displayed in Supplementary Figure 1. If participants answered the questionnaire twice, i.e. attended an assessment centre for the repeat visit, the answer at the second time point was used in analysis, in order to increase the mean age of the sample. To reduce the likelihood of including congenital forms of deafness, participants who selected ‘I am completely deaf’ in the UKBB questionnaire were excluded from analysis.

Note that a further, objective measure of hearing, the speech reception threshold using the ‘Digits in Noise’ (DIN) protocol, was obtained from 160,955 of the UK Biobank participants^45,46^ Preliminary heritability assessment of the DIN did not yield clear heritability or association with age and therefore it was not considered suitable for the present study.

Questionnaire responses for the ELSA and TwinsUK replication samples were derived to obtain comparable phenotypes to the UKBB phenotype (Supplementary Figure 1). For the ELSA sample, case/control phenotypes were derived from responses to questionnaire measures collected during study Wave 7. The *HDiff* phenotype was derived using responses from two questions; “Do you ever have any difficulties with your hearing?” and “Do you find it difficult to follow a conversation if there is background noise, such as TV, radio or children playing (using a hearing aid as usual)?” Cases consist of participants who responded “Yes” to both questions, and controls who responded “No” to both questions. As in the UKBB analysis, controls who report hearing aid use or age <50 were removed, as were any cochlear implant users in the case or control samples. The *HAid* phenotype was derived from responses to the question “Whether ever wears a hearing aid”; cases responded “Yes most of the time”, or “Yes some of the time” while controls responded “No”. During ELSA data processing, age is capped at 90 years, and thus individuals aged > 90 are reported to be 90 years of age. Association analysis *HDiff* ELSA sample N = 3545 and *HAid* ELSA sample N = 4482.

The TwinsUK phenotypes were likewise derived from responses to questionnaire measures. *HDiff* cases responded either “Yes, diagnosed by doctor or health professional” or “Yes, not diagnosed by health professional” to the question “Do you suffer from hearing loss?” while controls responded “No”. *HAid* cases responded or indicated “Yes” to either of “Do you wear a hearing aid?” and ‘Wearing a hearing aid’. *HAid* controls responded “No”. As TwinsUK is a longitudinal study, a number of participants gave responses to the same questions on multiple occasions. The most recent response was included in analysis, unless the latest response indicated that hearing had improved. In this scenario, the participant was excluded. Twins aged <40 were removed from analysis. Association analysis *HDiff* TwinsUK sample N = 3636 and *HAid* TwinsUK sample N = 3435.

### Genotyping and imputation

The ~500,000 samples in UKBB were genotyped on one of two arrays; 50,000 samples were genotyped on the Affymetrix UK BiLEVE Axiom array while the remaining ~450,000 were genotyped on the Affymetrix UK Biobank Axiom® array. The two arrays shared 95% coverage resulting in >800,000 genotyped SNPs. Imputation was carried out centrally by UKBB, primarily using the HRC reference panel and IMPUTE2^47^. SNPs which do not feature on this panel were imputed with the UK 10K and 1000G panel. Analysis in this study was conducted with version 3 of the UKBB imputed data with 487,409 samples imputed and available for analysis following UKBB centrally performed QC filters.

ELSA samples were genotyped at UCL Genomics in two batches using the Illumina HumanOmni 2.5M platform. Imputation was carried out centrally by ELSA with IMPUTE2, using the 1000 Genomes phase I data set^48^ (https://www.elsa-project.ac.uk/uploads/elsa/elsa_analysis.pdf).

Genotyping of TwinsUK was conducted with a combination of Illumina arrays; HumanHap300, HumanHap610Q, 1M-Duo and 1.2MDuo 1M. The imputation reference was 1000G Phase3 v5 (GRCh37).

### Statistical analysis

Discovery association was performed using a linear mixed-effects model approach to test for association between imputed SNP dosages and the two traits. BOLT-LMM v.2^25^ was used for the association analysis, which corrects for population stratification and within-sample relatedness. In addition, the analysis was adjusted for age, sex, UKBB genotyping platform and UKBB PCs1-10. For quality control, SNPs were filtered based on two thresholds: (1) minor allele frequency (MAF) ≥ 0.01; and (2) INFO score > 0.7. By implementing an MAF cutoff of 0.01, we reduced the likelihood of including participants with forms of congenital deafness, as we only detected variants that occur at least in 1/100 participants, a higher rate of variants than the rate of congenital deafness. Individuals with < 98% genotype call rate were removed. Conditional and joint SNP analysis was performed to identify independent signals within highly associated regions, using GCTA-COJO^26^. This analysis requires the linkage disequilibrium reference sample, which was obtained by random selection of 10,000 individuals from the UKBB cohort with White British ancestry. The reference sample size was selected to maximise power based on previous data simulations^49^. Independent SNPs identified with GCTA-COJO were mapped to the nearest protein coding gene using variant effect predictor (VEP), genome build GRCh37. VEP was used to establish whether the SNP was in an exonic, intronic or intergenic region, and also the functional consequence of the variant at that position. Univariate linkage disequilibrium (LD) score regression was used to calculate whether inflated test statistics were likely due to the polygenic nature of the trait or confounding bias, by analysing the relationship between test statistic and LD^50^.

SNP heritability estimates for the two traits were calculated with BOLT-LMM (h2g) and recalculated to the liability scale, with sample and population prevalence as per the case prevalence in the analysed sample; *HDiff* at 0.35 and *HAid* at 0.052.

SNPs identified with conditional analysis (Table 1) were tested for association with *HDiff* and *HAid* phenotypes in each of the three cohorts UKBB (non-white British), TwinsUK and ELSA. The UKBB white non-British sample was examined using the same protocol as the White British dataset described above, under the linear mixed models method with BOLT-LMM adjusting for age, sex, UKBB PCs 1-10 and genotyping platform. The TwinsUK sample was analysed using a linear mixed-effects model regression adjusting for age and sex with GEMMA^51^, accoutning for family structure. The ELSA samples for *HDiff* and *HAid* are <5,000 and one of each pair of related individuals was excluded from analysis (relatedness was estimated in PLINK 1.9^52^), therefore PLINK2 logistic regression was used to test for association in the ELSA sample, adjusting for age and sex.

For SNPs significantly associated with ARHI in the discovery, a fixed-effect inverse-variance weighted meta-analysis was conducted using METAL^53^ version 2011-03-25 with the three samples: white non-British UKBB, ELSA and TwinsUK. BOLT-LMM does not report analysed sample size per SNP, so to obtain the weight of the UKBB replication sample per SNP, sample size was calculated from PLINK linear regression analysis.

### Gene prioritization, pathway and tissue enrichment analysis

Summary statistics from the UKBB *HDiff* trait were input for Functional Mapping and Annotation of Genome-wide Association Studies (FUMA)^54^ as an alternative way to identify independent significant SNPs, lead SNPs, and functional annotations. Firstly, SNP2GENE function within FUMA was used to identify (i) *independent significant SNPs* (P≤5E-08) that were independent from each other at r^2^<0.6, and (ii) *lead SNPs -* significant SNPs that were independent from each other at r^2^<0.1. In addition, genomic risk loci borders were determined using candidate/tagged SNPs, which were SNPs in LD with independent significant SNPs at P ≤ 5E-08 and r^2^ ≥ 0.6. Secondly, lead SNPs were mapped to the nearest protein coding genes with a maximum distance of 10kb using VEP^27^. Gene set enrichment analysis was performed using ToppGene Suite^55^. These two steps were repeated with a genome-wide suggestive level (P ≤ 1E-05) to highlight regions that were significant and suggestive of harbouring causal variants. Alongside SNP-based analysis, we analysed the hearing difficulty GWAS using MAGMA^34^, a gene-based method which has been made available within FUMA. In MAGMA, the effect of multiple SNPs is combined together by mapping SNPs to 19,146 protein coding genes based on genomic location of 10kb to the genes, and a P-value describing the association found with hearing difficulty was derived.

### Protein localisation in mouse tissue sections

Adult mouse cochleae were collected at p28-p30 from C57BL/6 mice, bred in an in-house facility. Mice were euthanised according to Schedule 1 procedures as described in United Kingdom legislation outlined in the Animals (Scientific Procedures) Act 1986. Dissected inner ears were fixed in 4% paraformaldehyde diluted in PBS for 1 hour at room temperature before being washed several times in PBS. They were then decalcified in 10% EDTA overnight at 4°C, before being separated from the vestibular system. Cochlea were mounted in 4% low-melting point agarose and sectioned on a Vibratome (1000 plus system, Intracel) at 200-µm intervals. Antibodies used to identify protein localisation in the organ of Corti were: nidogen-2 (NID2) at 1:750 dilution (Ab14513, Abcam), clarin-2 (CLRN2) at 1:1000 (HPA042407, Atlas Antibodies) and rho guanine nucleotide exchange factor 28 (ARHGEF28) at 1:1000 (HPA037602, Atlas Antibodies). All were detected using of an isotype-specific secondary antibody, Alexa Fluor 488 goat anti-rabbit (Santa Cruz Biotechnology). Antibodies were diluted in a goat blocking solution (4% triton, 8% goat serum, 1g BSA, 50ml PHEM buffer) and sections were stained with primary antibodies overnight at 4°C. Following PBS washes, sections were incubated with the secondary antibody at 1:1000 in darkness at room temperature for 2 hours. Phalloidin-Atto 647N to f-actin (Sigma-Aldrich, Gillingham, UK) and DAPI were added to the secondary antibody incubations at 1:1000 to stain hair cell stereocilia and DNA respectively. Samples were imaged using a Zeiss LSM 880 Airyscan 20x objective.

## Acknowledgments

The research was carried out using the UK Biobank Resource under application number 11516. HRRW is funded by a PhD Studentship Grant, S44, from Action on Hearing Loss. The study was also supported by funding from NIHR UCLH BRC Deafness and Hearing Problems Theme, MED_EL and the NIHR Manchester Biomedical Research Centre. The English Longitudinal Study of Ageing is jointly run by University College London, Institute for Fiscal Studies, University of Manchester and National Centre for Social Research. Genetic analyses have been carried out by UCL Genomics and funded by the Economic and Social Research Council and the National Institute on Aging. All GWAS data have been deposited in the European Genome-phenome Archive. Data governance was provided by the METADAC data access committee, funded by ESRC, Wellcome, and MRC. (2015-2018: Grant Number MR/N01104X/1 2018-2020: Grant Number ES/S008349/1). TwinsUK is funded by the Wellcome Trust, Medical Research Council, European Union, the National Institute for Health Research (NIHR)-funded BioResource, Clinical Research Facility, and Biomedical Research Centre based at Guy’s and St Thomas’ NHS Foundation Trust in partnership with King’s College London. We would like to thank all the participants of UK Biobank, English Longitudinal Study of Aging and TwinsUK.

## Author contributions

Helena RR. Wells, Maxim B. Freidin and Fatin N. Zainul Abidin performed the analysis. Helena RR. Wells, Maxim B. Freidin, Frances MK. Williams and Sally J Dawson designed, wrote and oversaw the study. Antony Payton, Piers Dawes, Kevin J. Munro, Cynthia C. Morton and David R. Moore all contributed to the study concept and writing.

## Competing interests

David R. Moore, Scientific Advisor: hearX Ltd., Otonomy Inc

## Availability of data

Data that support the findings of this study are publically available upon successful application from the UK Biobank, the English Longitudinal Study of Aging and TwinsUK.

Derived data from the UK Biobank data fields and GWAS summary statistics that support the findings of this study, will be made available as part of the UK Biobank Returns Catalogue following the publication of this manuscript.

## References

1. WHO | Estimates. Who (2018). Retrieved from www.who.int/pbd/deafness/estimates/en/

2. Bedin, E. et al. Age-related hearing loss in four Italian genetic isolates: An epidemiological study. Int. J. Audiol. 48, 465–472 (2009).

3. Bogo, R. et al. The Role of Genetic Factors for Hearing Deterioration Across 20 Years: A Twin Study. Journals Gerontol. Ser. A Biol. Sci. Med. Sci. 70, 647–653 (2015).

4. Gates, G. A., Couropmitree, N. N. & Myers, R. H. Genetic Associations in Age-Related Hearing Thresholds. Arch. Otolaryngol. Neck Surg. 125, 654 (1999).

5. Sudlow, C. et al. UK Biobank: An Open Access Resource for Identifying the Causes of a Wide Range of Complex Diseases of Middle and Old Age. PLOS Med. 12, e1001779 (2015).

6. Brewster, K. K. et al. Age-Related Hearing Loss and Its Association with Depression in Later Life. Am. J. Geriatr. Psychiatry 26, 788–796 (2018).

7. Rutherford, B. R., Brewster, K., Golub, J. S., Kim, A. H. & Roose, S. P. Sensation and Psychiatry: Linking Age-Related Hearing Loss to Late-Life Depression and Cognitive Decline. Am. J. Psychiatry 175, 215–224 (2018).

8. Han, J. H., Lee, H. J., Jung, J. & Park, E.-C. Effects of self-reported hearing or vision impairment on depressive symptoms: a population-based longitudinal study. Epidemiol. Psychiatr. Sci. 1–13 (2018). doi:10.1017/S2045796018000045

9. Jayakody, D. M. P., Friedland, P. L., Martins, R. N. & Sohrabi, H. R. Impact of Aging on the Auditory System and Related Cognitive Functions: A Narrative Review. Front. Neurosci. 12, 125 (2018).

10. Wei, J. et al. Hearing Impairment, Mild Cognitive Impairment, and Dementia: A Meta-Analysis of Cohort Studies. Dement. Geriatr. Cogn. Dis. Extra 7, 440–452 (2017).

11. Gurgel, R. K. et al. Relationship of hearing loss and dementia: a prospective, population-based study. Otol. Neurotol. 35, 775–81 (2014).

12. Lin, F. R. et al. Hearing loss and incident dementia. Arch. Neurol. 68, 214–20 (2011).

13. Vos, T. et al. Global, regional, and national incidence, prevalence, and years lived with disability for 301 acute and chronic diseases and injuries in 188 countries, 1990-2013: a systematic analysis for the Global Burden of Disease Study 2013. The Lancet 386, (2015).

14. Fransen, E. et al. Genome-wide association analysis demonstrates the highly polygenic character of age-related hearing impairment. Eur. J. Hum. Genet. 23, 110–115 (2015).

15. Newman, D. L. et al. GRM7 variants associated with age-related hearing loss based on auditory perception. Hear. Res. 294, 125–132 (2012).

16. Nolan, L. S. et al. Estrogen-related receptor gamma and hearing function: evidence of a role in humans and mice. Neurobiol. Aging 34, 2077.e1-9 (2013).

17. Van Laer, L. et al. A genome-wide association study for age-related hearing impairment in the Saami. Eur. J. Hum. Genet. 18, 685–693 (2010).

18. Luo, H. et al. Association of GRM7 Variants with Different Phenotype Patterns of Age-Related Hearing Impairment in an Elderly Male Han Chinese Population. PLoS One 8, e77153 (2013).

19. Friedman, R. A. et al. GRM7 variants confer susceptibility to age-related hearing impairment. Hum. Mol. Genet. 18, 785–796 (2009).

20. Duijvestijn, J. A., Anteunis, L. J., Hendriks, J. J. & Manni, J. J. Definition of hearing impairment and its effect on prevalence figures. A survey among senior citizens. Acta Otolaryngol. 119, 420–3 (1999).

21. Wolber, L. E. et al. Salt-inducible kinase 3, SIK3, is a new gene associated with hearing. Hum. Mol. Genet. 23, 6407–18 (2014).

22. Vuckovic, D. et al. Genome-wide association analysis on normal hearing function identifies PCDH20 and SLC28A3 as candidates for hearing function and loss. Hum. Mol. Genet. 24, 5655–64 (2015).

23. Girotto, G. et al. Hearing function and thresholds: a genome-wide association study in European isolated populations identifies new loci and pathways. J. Med. Genet. 48, 369–74 (2011).

24. Hoffmann, T. J. et al. A Large Genome-Wide Association Study of Age-Related Hearing Impairment Using Electronic Health Records. PLoS Genet. 12, e1006371 (2016).

25. Loh, P.-R. et al. Efficient Bayesian mixed-model analysis increases association power in large cohorts. Nat. Genet. 47, 284–90 (2015).

26. Yang, J. et al. Conditional and joint multiple-SNP analysis of GWAS summary statistics identifies additional variants influencing complex traits. Nat. Genet. 44, 369–75, S1-3 (2012).

27. McLaren, W. et al. The Ensembl Variant Effect Predictor. Genome Biol. 17, 122 (2016).

28. Schönberger, J. et al. Mutation in the transcriptional coactivator EYA4 causes dilated cardiomyopathy and sensorineural hearing loss. Nat. Genet. 37, 418–422 (2005).

29. Makishima, T. et al. Nonsyndromic hearing loss DFNA10 and a novel mutation of *EYA4* : Evidence for correlation of normal cardiac phenotype with truncating mutations of the Eya domain. Am. J. Med. Genet. Part A 143A, 1592–1598 (2007).

30. Pfister, M. et al. A 4-bp insertion in the eya-homologous region (eyaHR) of EYA4 causes hearing impairment in a Hungarian family linked to DFNA10. Mol. Med. 8, 607–11 (2002).

31. Sheets, L., Trapani, J. G., Mo, W., Obholzer, N. & Nicolson, T. Ribeye is required for presynaptic Ca(V)1.3a channel localization and afferent innervation of sensory hair cells. Development 138, 1309–19 (2011).

32. Shahin, H. et al. Mutations in a novel isoform of TRIOBP that encodes a filamentous-actin binding protein are responsible for DFNB28 recessive nonsyndromic hearing loss. Am. J. Hum. Genet. 78, 144–52 (2006).

33. Riazuddin, S. et al. Mutations in TRIOBP, which encodes a putative cytoskeletal-organizing protein, are associated with nonsyndromic recessive deafness. Am. J. Hum. Genet. 78, 137–43 (2006).

34. de Leeuw, C. A., Mooij, J. M., Heskes, T. & Posthuma, D. MAGMA: Generalized Gene-Set Analysis of GWAS Data. PLOS Comput. Biol. 11, e1004219 (2015).

35. Fields, R. R. et al. Usher Syndrome Type III: Revised Genomic Structure of the USH3 Gene and Identification of Novel Mutations. Am. J. Hum. Genet. 71, 607–617 (2002).

36. Adato, A. et al. USH3A transcripts encode clarin-1, a four-transmembrane-domain protein with a possible role in sensory synapses. Eur. J. Hum. Genet. 10, 339–350 (2002).

37. Kohfeldt, E., Sasaki, T., Göhring, W. & Timpl, R. Nidogen-2: a new basement membrane protein with diverse binding properties. J. Mol. Biol. 282, 99–109 (1998).

38. Cañete-Soler, R., Wu, J., Zhai, J., Shamim, M. & Schlaepfer, W. W. p190RhoGEF Binds to a destabilizing element in the 3’ untranslated region of light neurofilament subunit mRNA and alters the stability of the transcript. J. Biol. Chem. 276, 32046–50 (2001).

39. Volkening, K., Leystra-Lantz, C. & Strong, M. J. Human low molecular weight neurofilament (NFL) mRNA interacts with a predicted p190RhoGEF homologue (RGNEF) in humans. Amyotroph. Lateral Scler. 11, 97–103 (2010).

40. Rico, B. et al. Control of axonal branching and synapse formation by focal adhesion kinase. Nat. Neurosci. 7, 1059–1069 (2004).

41. Droppelmann, C. A. et al. Detection of a novel frameshift mutation and regions with homozygosis within ARHGEF28 gene in familial amyotrophic lateral sclerosis. Amyotroph. Lateral Scler. Front. Degener. 14, 444–451 (2013).

42. Bycroft, C. et al. Genome-wide genetic data on ~500,000 UK Biobank participants. bioRxiv 166298 (2017). doi:10.1101/166298

43. Steptoe, A., Breeze, E., Banks, J. & Nazroo, J. Cohort profile: the English longitudinal study of ageing. Int. J. Epidemiol. 42, 1640–8 (2013).

44. Moayyeri, A., Hammond, C. J., Hart, D. J. & Spector, T. D. The UK Adult Twin Registry (TwinsUK Resource) Europe PMC Funders Group. Twin Res Hum Genet 16, 144–149 (2013).

45. Dawes, P. et al. Hearing in middle age: a population snapshot of 40-to 69-year olds in the United Kingdom. Ear Hear. 35, e44–51 (2014).

46. Moore, D. R. et al. Relation between speech-in-noise threshold, hearing loss and cognition from 40-69 years of age. PLoS One 9, e107720 (2014).

47. Howie, B. N., Donnelly, P. & Marchini, J. A Flexible and Accurate Genotype Imputation Method for the Next Generation of Genome-Wide Association Studies. PLoS Genet. 5, e1000529 (2009).

48. Howie, B., Fuchsberger, C., Stephens, M., Marchini, J. & Abecasis, G. R. Fast and accurate genotype imputation in genome-wide association studies through pre-phasing. Nat. Genet. 44, 955–959 (2012).

49. Yang, J. et al. Conditional and joint multiple-SNP analysis of GWAS summary statistics identifies additional variants influencing complex traits. Nat. Genet. 44, 369–375 (2012).

50. Zheng, J. et al. LD Hub: a centralized database and web interface to perform LD score regression that maximizes the potential of summary level GWAS data for SNP heritability and genetic correlation analysis. Bioinformatics 33, 272–279 (2017).

51. Zhou, X. & Stephens, M. Genome-wide efficient mixed-model analysis for association studies. Nat. Genet. 44, 821–4 (2012).

52. Chang, C. C. et al. Second-generation PLINK: rising to the challenge of larger and richer datasets. Gigascience 4, 7 (2015).

53. Willer, C. J., Li, Y. & Abecasis, G. R. METAL: fast and efficient meta-analysis of genomewide association scans. Bioinforma. Appl. NOTE 26, 2190–2191 (2010).

54. Watanabe, K., Taskesen, E., van Bochoven, A. & Posthuma, D. Functional mapping and annotation of genetic associations with FUMA. Nat. Commun. 8, 1826 (2017).

55. Chen, J., Bardes, E. E., Aronow, B. J. & Jegga, A. G. ToppGene Suite for gene list enrichment analysis and candidate gene prioritization. Nucleic Acids Res. 37, W305–11 (2009).

