## Supplementary Tables … Figures for "Genome-wide association study identifies 44 independent genomic loci for self-reported adult hearing difficulty in the UK Biobank cohort"

### Supplementary Figures & Tables

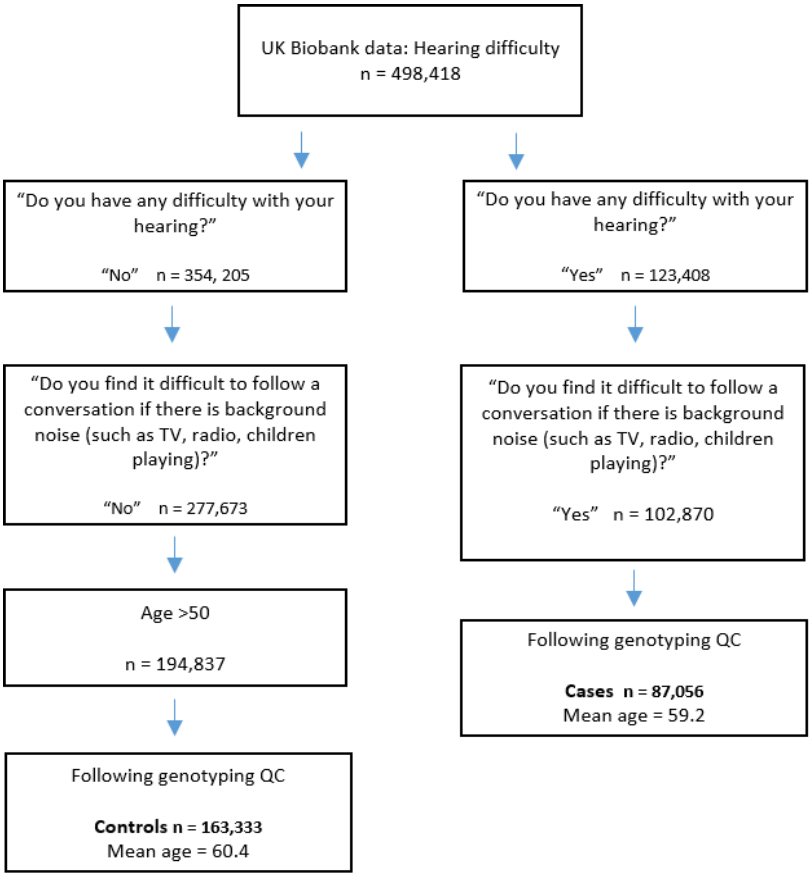


**Supplementary Figure 1a. Flow chart describing case-control assignment for the hearing difficulty (HDiff) phenotype.** Participants answered questions as part of the UKBB questionnaire administered at UKBB assessment centres. Participants who answered 'Prefer not to answer', 'I am completely deaf' and 'Do not know' were removed from the analysis. Participants were removed from the control group if they answered "Yes" to "Do you use a hearing aid most of the time?" A lower age limit of 50 was implemented for controls to ensure age was consistent between the case and control groups due to the association of aging with the trait.

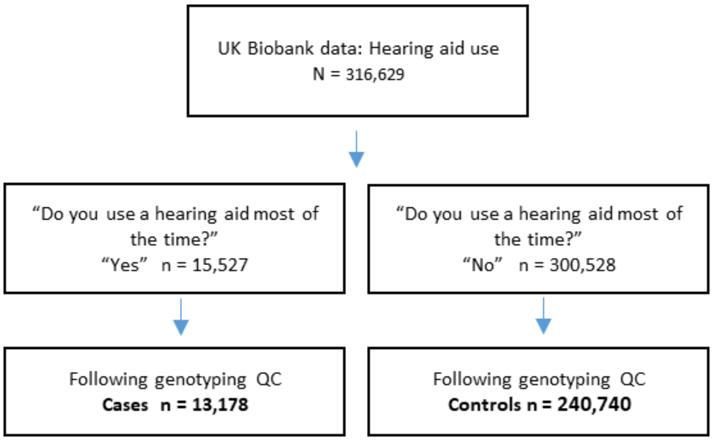


**Supplementary Figure 1b. Flow chart describing case-control assignment for the hearing aid use (*HAid*) phenotype**. Participants answered questions as part of the UKBB questionnaire administered at the UKBB assessment centres. No information was collected regarding age at hearing aid prescription or cause of hearing loss.

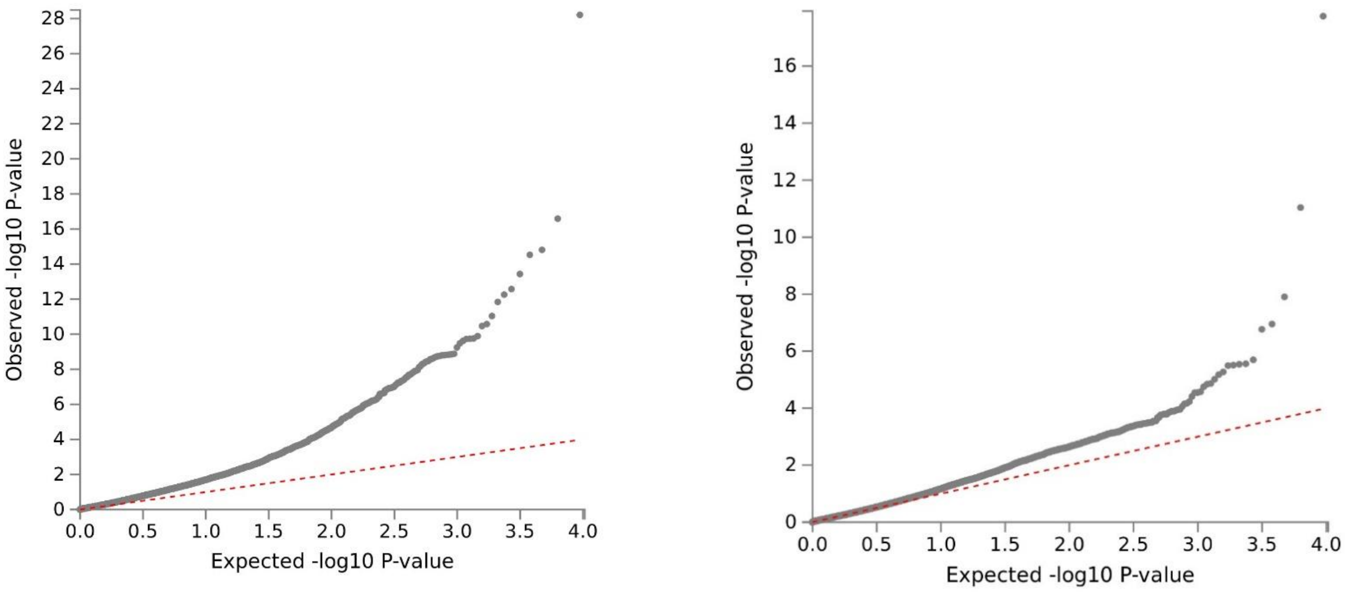


**Supplementary Figure 2a. Q-Q plot of GWAS summary statistics, HDiff (left) and HAid (right).**

Univariate linkage disequilibrium score (LDSC) regression estimated the polygenic nature of the HDiff trait caused 8% of the inflation; the intercept was 1.032. For HAid, the ratio was estimated at 5% with an intercept of 1.03.

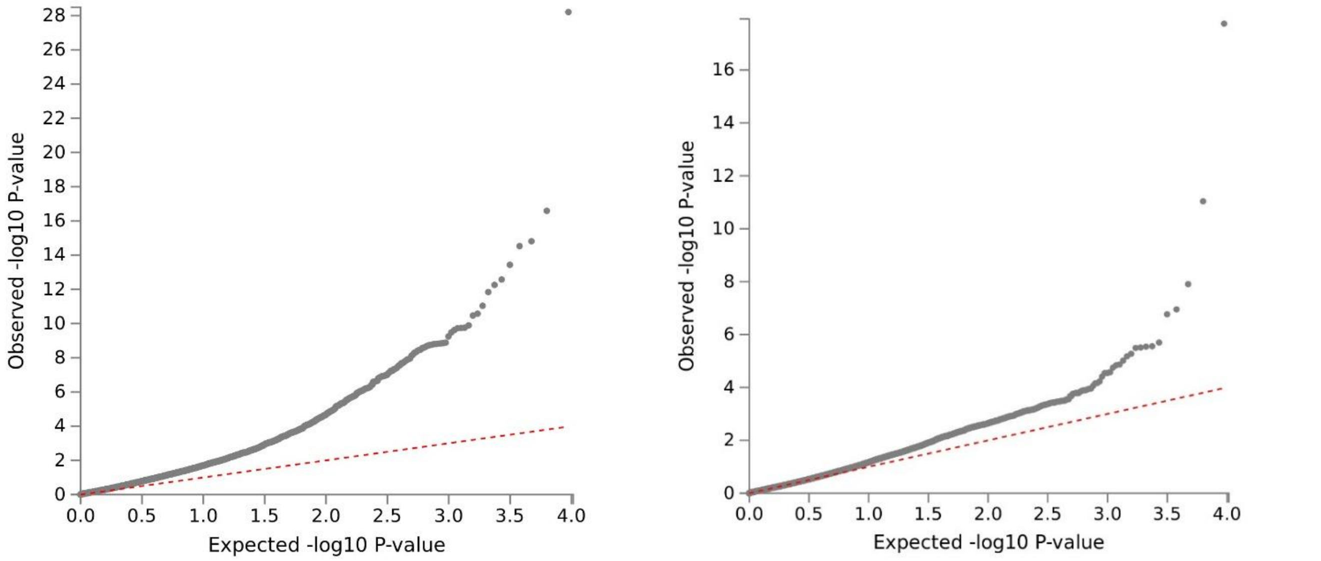


**Supplementary Figure 2b. Q-Q plot of the gene-based test computed by MAGMA, HDiff (left) and HAid (right).**

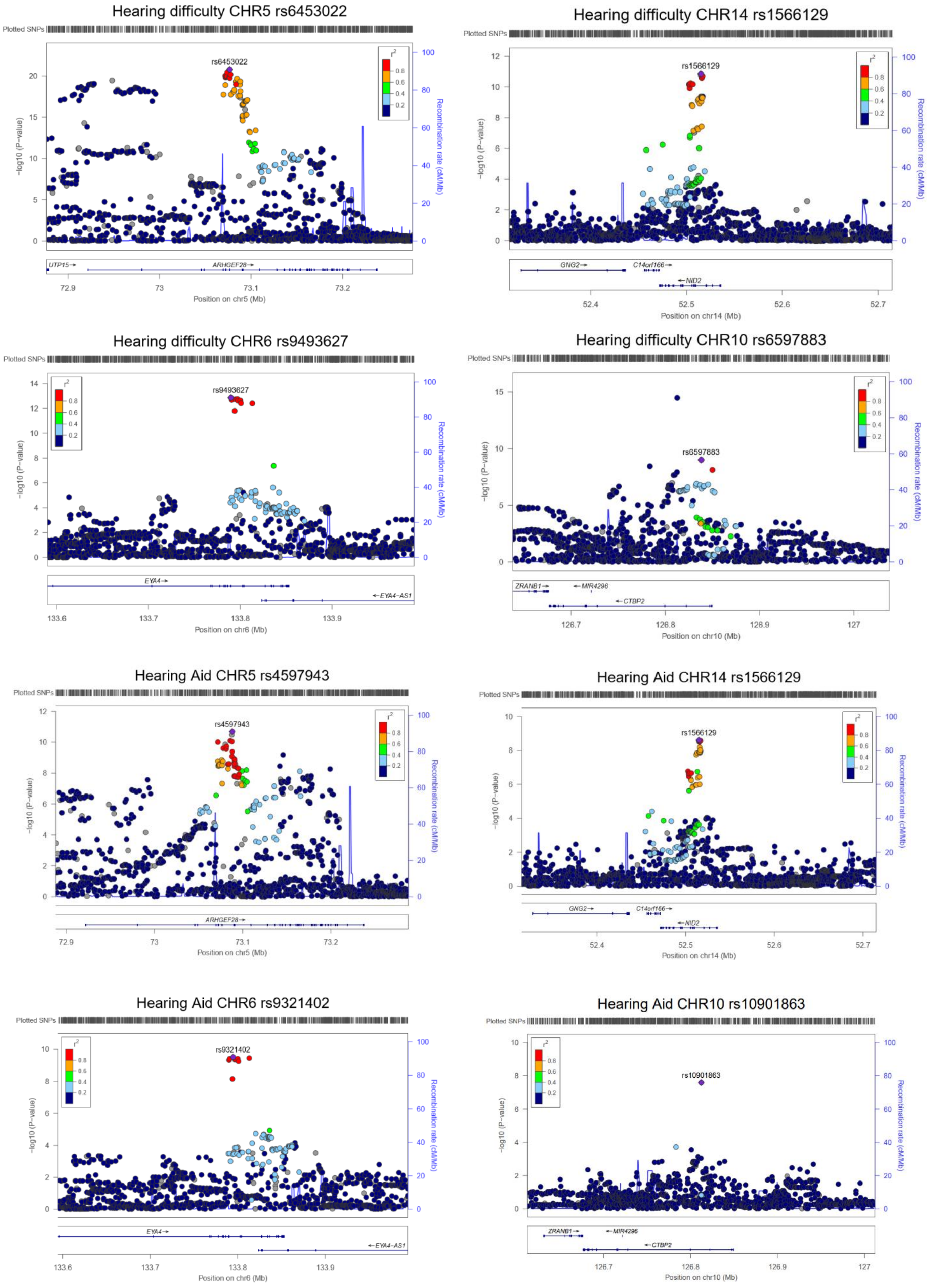
Hearing difficulty CHR5 rs6453022
rs6453022
UTP15
ARHGEF28
Position on chr5 (Mb)
Hearing difficulty CHR14 rs1566129
rs1566129
GNG2
C14orf166
NID2
Position on chr14 (Mb)
Hearing difficulty CHR6 rs9493627
rs9493627
EYA4
EYA4-AS1
Position on chr6 (Mb)
Hearing difficulty CHR10 rs6597883
rs6597883
ZRANB1
MIR4296
CTBP2
Position on chr10 (Mb)
Hearing Aid CHR5 rs4597943
rs4597943
ARHGEF28
Position on chr5 (Mb)
Hearing Aid CHR14 rs1566129
rs1566129
GNG2
C14orf166
NID2
Position on chr14 (Mb)
Hearing Aid CHR6 rs9321402
rs9321402
EYA4
EYA4-AS1
Position on chr6 (Mb)
Hearing Aid CHR10 rs10901863
rs10901863
ZRANB1
MIR4296
CTBP2
Position on chr10 (Mb)
Plotted SNPs
-log10 (P-value)
Recombination rate (cM/Mb)

**Supplementary Figure 3. Locus plots displaying regions significantly associated in both *HDiff* and *HAid* analysis.** Locus plots generated with HDiff summary statistics. Purple indicates lead independent SNP generated from GCTA-COJO conditional analysis. The colouring of remaining SNPs represents the correlation ($r^2$) to the lead SNP (purple).

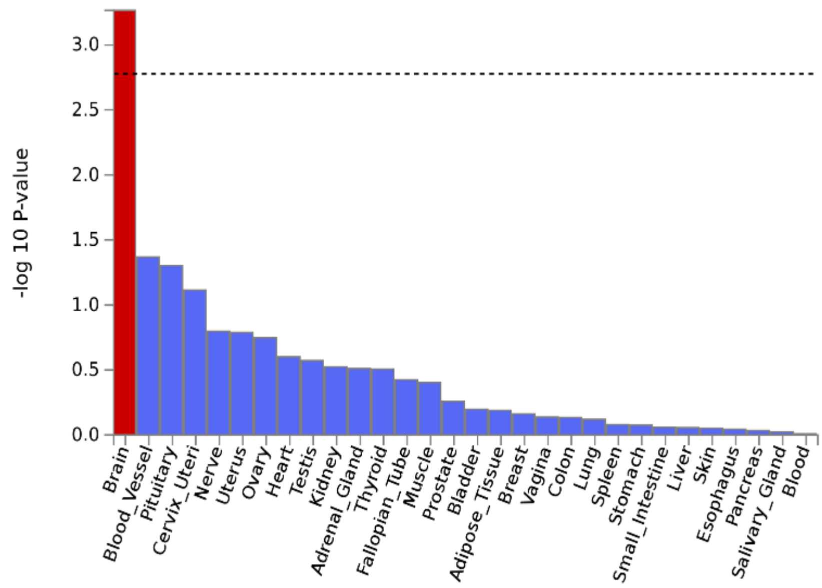


**Supplementary Figure 4. MAGMA gene-property analysis for tissue specificity from average expression of 30 general tissue types from GTEx v6.** Relationships between tissue specific gene expression profiles (x-axis) and genetic association of genes (y-axis) is shown. Genetic associations of genes were performed in MAGMA and represent the aggregate effect of all SNPs in a gene. The dotted line indicates the Bonferroni-corrected level of 2.6E-06.

| Supplementary Table 1. Summary statistics for *HDiff* phenotype from the replication meta-analysis of the white non-British UKBB sample, TwinsUK and ELSA | | | | | | |
|---|---|---|---|---|---|---|
| Marker Name | **Allele1** | **Allele2** | **Weight** | **Zscore** | **P-value** | **Direction** |
| rs759016271 | a | agtagtccacttttcttctttgcctg | 29866 | 4.556 | 5.20E-06 | +++ |
| rs1566129 | t | c | 30894 | 3.534 | 0.00041 | +++ |
| rs36062310 | a | g | 30868 | 2.974 | 0.002938 | +-- |
| rs143282422 | a | g | 30274 | 2.652 | 0.007996 | +-+ |
| rs12225399 | c | g | 29802 | 2.624 | 0.008688 | +++ |
| rs6597883 | t | c | 30274 | 2.585 | 0.009748 | ++- |
| rs62033400 | a | g | 30851 | 2.513 | 0.01198 | +++ |
| rs7951935 | t | g | 29802 | 2.48 | 0.01312 | ++- |
| rs141403654 | a | t | 29802 | -2.392 | 0.01674 | --- |
| rs35186928 | a | g | 29866 | 2.348 | 0.01885 | +-+ |
| rs17671352 | t | c | 30852 | 2.228 | 0.02587 | ++- |
| rs217289 | a | g | 29866 | 2.227 | 0.02597 | +-+ |
| rs62188635 | t | c | 30377 | -2.189 | 0.02859 | --- |
| rs76837345 | a | g | 30318 | -2.158 | 0.0309 | --- |
| rs55635402 | a | g | 29802 | 2.132 | 0.03301 | +++ |
| rs10824108 | t | g | 30274 | 1.982 | 0.04752 | +-+ |
| rs2236401 | t | c | 29866 | 1.873 | 0.06102 | ++- |
| rs5756795 | t | c | 30868 | -1.696 | 0.08981 | --+ |
| rs7525101 | t | c | 30389 | 1.636 | 0.1018 | +++ |
| rs10475169 | a | c | 30356 | -1.603 | 0.1088 | -+- |
| rs4948502 | t | c | 30274 | 1.574 | 0.1155 | +-+ |
| rs9691831 | a | g | 30717 | -1.512 | 0.1305 | -+- |
| rs12938775 | a | g | 30852 | -1.481 | 0.1385 | -+- |
| rs4611552 | t | c | 30753 | -1.434 | 0.1515 | --- |
| rs13093972 | a | g | 30098 | -1.413 | 0.1577 | --+ |
| rs6453022 | a | c | 30356 | 1.327 | 0.1846 | +-- |
| rs835267 | a | g | 30274 | 1.17 | 0.242 | ++- |
| rs35414371 | a | t | 30856 | 1.163 | 0.245 | +-+ |
| rs9493627 | a | g | 29866 | 1.018 | 0.3086 | +++ |
| rs12027345 | a | g | 30389 | -0.986 | 0.3244 | --+ |
| rs3890736 | a | g | 30318 | -0.706 | 0.4805 | +-- |
| rs4947828 | t | g | 30717 | -0.562 | 0.574 | +-- |

| | | | | | | |
|---|---|---|---|---|---|---|
| rs13277721 | a | g | 30318 | 0.527 | 0.5983 | +-- |
| rs12552 | a | g | 30676 | 0.489 | 0.6249 | +-+ |
| rs6890164 | a | g | 30356 | 0.298 | 0.7658 | +-- |
| 3:182069497_TA_T | t | ta | 30098 | 0.298 | 0.766 | ++- |
| rs132929 | a | g | 30868 | 0.234 | 0.815 | ++- |
| rs10927035 | t | c | 30389 | 0.219 | 0.8268 | +-+ |
| rs34442808 | t | ta | 30356 | 0.218 | 0.8277 | -++ |
| rs9366417 | a | g | 29866 | 0.137 | 0.8911 | +-+ |
| rs62015206 | t | c | 30459 | -0.071 | 0.9437 | --+ |

**Supplementary Table 1.** Marker Name, SNP ID; Allele1, the first allele for this marker in the first file where it occurs; Allele 2, the second allele for this marker in the first file where it occurs; Weight, the sum of the individual study weights (N) for this marker; Z-score, the combined z-statistic for the marker; P-value, meta-analysis p-value; Direction, direction of effect for each study ordered white non-British UKBB sample, TwinsUK, ELSA.

| Supplementary Table 2. Summary statistics for the *HAid* phenotype in the replication meta-analysis of white non-British UKBB sample, TwinsUK and ELSA | | | | | | |
|---|---|---|---|---|---|---|
| **Marker Name** | **Allele1** | **Allele2** | **Weight** | **Zscore** | **P-value** | **Direction** |
| rs4597943 | t | g | 34475 | 2.833 | 0.004608 | +++ |
| rs7823971 | a | c | 34919 | -1.886 | 0.05934 | --- |
| rs3915060 | t | c | 34359 | -1.664 | 0.09605 | --+ |
| rs1566129 | t | c | 35139 | 0.784 | 0.4333 | ++- |
| rs10901863 | t | c | 32251 | 0.638 | 0.5234 | +-- |
| rs9321402 | a | g | 35101 | 0.58 | 0.5622 | ++- |
| rs9677089 | a | c | 34727 | -0.537 | 0.5915 | --+ |

**Supplementary Table 2.** Marker Name, SNP ID; Allele1, the first allele for this marker in the first file where it occurs; Allele 2, the second allele for this marker in the first file where it occurs; Weight, the sum of the individual study weights (N) for this marker; Z-score, the combined z-statistic for the marker; P-value, meta-analysis p-value; Direction, direction of effect for each study ordered white non-British UKBB sample, TwinsUK, ELSA.

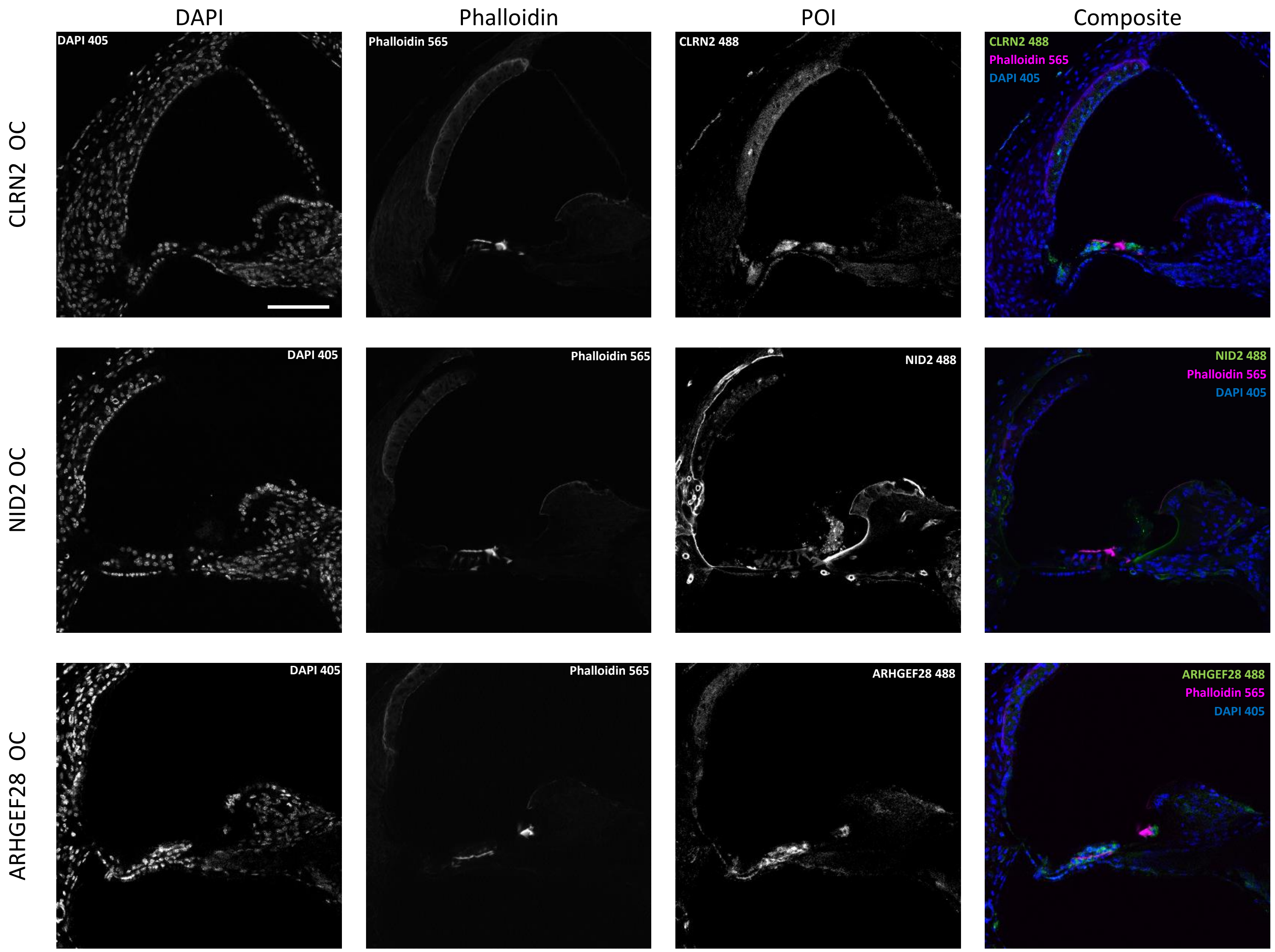
DAPI
Phalloidin
POI
Composite
CLRN2 OC
NID2 OC
ARHGEF28 OC
DAPI 405
Phalloidin 565
CLRN2 488
NID2 488
ARHGEF28 488

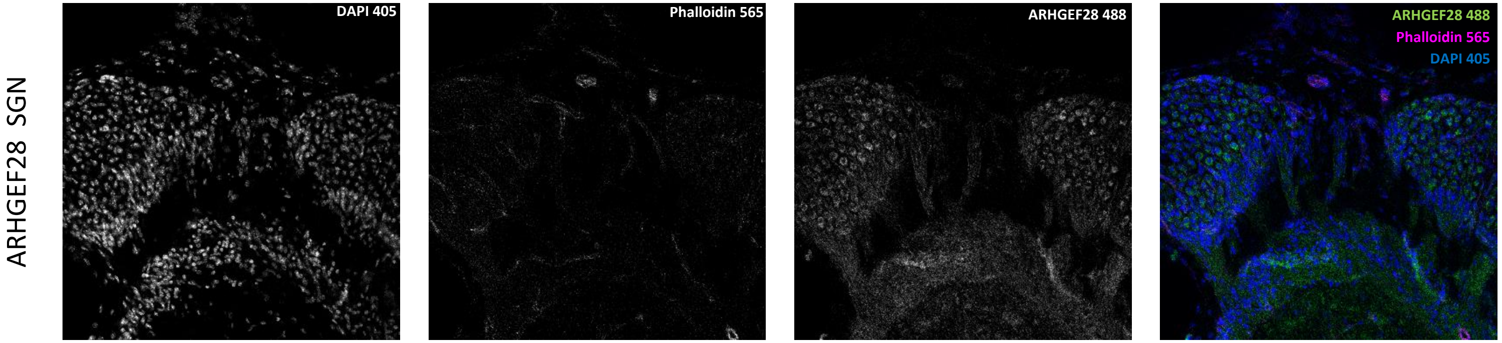


**Supplementary Figure 5. Immunofluorescence images of adult mouse cochlea Vibratome sections stained with proteins of interest (green), DAPI (blue) and Phalloidin (magenta)**

Individual channels presented in greyscale, and combined for composite colour images as in Figure 4. Three panels display the organ of Corti (OC) regions (CLRN2 OC, NID2 OC, and ARHGEF28 OC) and one displays the Spiral Ganglion Neurons (SGN) region (ARHGEF28 SGN).

The scale bar in the top left image displays 100µm. This scale is consistent for all images in this figure.
